# INS-fMRI: a novel method for mapping mesoscale connectome networks *in vivo* in nonhuman primates

**DOI:** 10.64898/2026.09.16.751612

**Authors:** Xiao Du, Jianbao Wang, Ying Zhang, Liang Zhu, Songping Yao, Dengfeng Zhou, Kenneth E. Schriver, Augix Guohua Xu, Anna Wang Roe

**Affiliations:** Department of Neurology of the Second Affiliated Hospital and Interdisciplinary Institute of Neuroscience and Technology, Zhejiang University School of Medicine, Hangzhou, China; Interdisciplinary Institute of Neuroscience and Technology, Key Laboratory for Biomedical Engineering of Ministry of Education, College of Biomedical Engineering and Instrument Science, Zhejiang University, Hangzhou, China; School of Brain Science and Brain Medicine and Interdisciplinary Institute of Neuroscience and Technology, Zhejiang University School of Medicine, Hangzhou, China; Division of Translational Neuroscience, Nathan Kline Institute of Translational Neuroscienc, Orangeburg, NY; Departments of Psychiatry and Neuroscience, NYU Grossman School of Medicine, New York, NY

## Abstract

Mapping brain connections is paramount in neuroscience. Lacking has been a method that can systematically map mesoscale (≤1mm) connections in the brain *in vivo* at brainwide scale. Mesoscale mapping is particularly important for NHPs in which information is represented in submillimeter clusters (e.g. cortical columns, striosomes, thalamic ‘rods’). Here, we present a method called INS-fMRI, which comprises optical stimulation of single sites in the brain using pulsed Infrared Neural Stimulation (INS), coupled with mapping of functionally connected sites in the brain using ultrahigh field fMRI. It is the only method available for mapping mesoscale connections rapidly (multiple networks mapped within single MRI sessions), systematically (allowing comparison of networks within single individuals), *in vivo* (no animal sacrifice needed), at whole brain scale, and without viral transfection. This method has broad applicability to a wide range of neuroscience questions. It can be used at any brain site, on different animal models, and on different MRI platforms. The ability to probe circuits repeatedly within single individuals also opens doors for studying changes in mesoscale circuits over time (e.g. for development, aging, disease studies). In addition, as this method circumvents the need for viral transfection, there is exciting potential for human clinical application.

## Introduction

### The problem

*Traditional approaches* to mapping brain circuits have relied on anatomical tracing. Such studies have formed the basis of much of our understanding of basic brain circuits and are considered ‘ground truth’ by which other studies are compared. The procedures typically involve injecting tracers and allowing time for transport (typically 2-3 weeks), followed by time-consuming reconstruction of labels in the brain ^1-3^. The number of tracers injected are limited to 3-4 tracers (injection sites), limiting the number of circuits that can be studied in a single animal. Study of multiple circuits requires use of many animals and also introduces the issue of inter-animal variability. In addition, the injections are often 1-5 mm in size, which, in primate brain, labels many submillimeter nodes of potentially different functionality, resulting in loss of connectional specificity and potentially incorrect inferences of connectivity.

*Need for new method*. We aimed to overcome some of these limitations by developing an ***in vivo* method** of mapping connections. Overcoming the need for animal sacrifice makes the study of many different circuits **within a single individual** possible, opening the doors for comparing the topography of different circuits, studying connectomes, node-hopping studies (stimulating connected sites), and developmental studies. A second requirement was to achieve **submillimeter precision**; this would permit the stimulation of a single functionally specific node (e.g. single orientation column) and study connections specific to this node. A third requirement was to enable study of **brainwide** connections, and not be limited to small fields of view. Finally, we envisioned a method that would be relatively **easy** to use, **rapid**, and **systematic**, making it broadly applicable.

*Organization of primate brains*. An important aspect of primate brains is the organization at submillimeter scale (cortical columns, striatal striosomes, thalamic rods). While focal tracer injections can label single columns, the small amount of tracer injected fails to effectively reveal distant, brainwide connections^4-6^. To achieve brainwide fields of view, magnetic resonance imaging (MRI) is needed. We therefore tested stimulation methods to focally activate single functionally specific, submillimeter nodes in primate cerebral cortex and mapped connected sites with optical imaging methods. Focal electrical^7-9,49-50^, optogenetic^10-12^, and infrared neural stimulation^13-15^ methods all demonstrated similar results that matched known anatomy, and strengthened the concept that, at local scale, cortical networks are based in columns^16^. However, these studies were limited to local (centimeter-scale) fields of view. Whether brainwide networks are based columnar connectivity remained unknown.

### Development of the protocol

*Previous methods*. There are many approaches to mapping connections in the brain. Each has its strengths and weaknesses. Resting state MRI^17,18^ is a non-invasive, whole brain scale method widely used in human studies. This method measures the correlation of BOLD signal in two areas of the brain, but interpretation of connectivity between areas is largely uncertain. With diffusion methods (DTI, DSI), it is difficult to achieve reliable connectivity with high resolution at whole brain scale^19^. Electrical stimulation comes with current spread making submillimeter stimulation difficult for brainwide scale studies and is less easy to conduct in the MRI environment (electrical-fMRI^20-22^). Optogenetics offers exciting cell type specificity. However, in the primate brain, while successfully conducted by a number of investigators (cf. optogenetic-fMRI^24,25^), viral transfection adds significant preparation time and procedures, is limited to the sites transfected, and can be can be unreliable^23^ .

### INS-fMRI

We developed a method called INS-fMRI (infrared neural stimulation with functional magnetic resonance imaging) to map brainwide circuits at millimeter resolution^26^. It has previously been shown that infrared light (wavelength 1875 nm, a peak of light absorption of water) can induce neuronal responses via induced transient temperature rise; with a specific pulsed paradigm (pulse train: 0.2msec per pulse, 200hz, for 0.5 sec), the absorption is confined to a very limited (submillimeter) volume^27^ (termed ‘thermal confinement’). Neuronal responses have been demonstrated via electrophysiological and 2-photon imaging methods^13,51^, as well as hemodynamic signatures of neuronal response, both optical imaging and fMRI, reflecting functional specificity, specificity of connectivity, and intensity dependence of stimulated and connected sites. INS can be applied safely without tissue damage in rat14, cat26, primate, and human cortex^14,26,47,53^, with a threshold of approximately 0.6J/cm^2^. Using a window of parameters, INS can be applied for 100s and 1000s of trials per day without tissue damage^14,47^, or apparent change in neural response when viewed with 2 photon imaging^51,52^ neural response, optical images, and fMRI response remain stable and modulation of circuitry and animal behavior remain stable. Based on Monte Carlo simulations and MR thermometry, temperature rise does not exceed 1°C, and rapidly dissipates due to the low duty cycle (0.2msec pulse per 5msec)^53,54^. Of the few mechanisms proposed for how temperature rise leads to neuronal response, in mammalian CNS, evidence suggests that heat transients induce changes in membrane capacitance and concomitant Na and K flux, leading to neural activation^28-^ 30.

### Challenges

The challenges of this development included MR engineering of multichannel RF coils to increase SNR^66^, calculation of SAR and conducting multiple studies to ensure our methods were non-damaging, development of spatial and temporal optical stimulation delivery methods, designing precision MR-compatible micromanipulators and advancers that could be controlled at a distance from the control room, development of identifying stimulation parameters that would be effective (lead to statistically robust and reproducible response at connected sites, and remain non-damaging) at brainwide scale, monitoring and maintainging physiological parameters of the monkey from a distance in the MRI, conducting data acquisition in an ultrahigh field MR scanner with minimal inhomogeneity, and developing analytical and statistical methods (e.g. removal or modification of standard steps in the fMRI analysis pipeline which induce blur^67^) to identify small but significant focal BOLD signals. To a large extent, these challenges have been met, as described below.

### Multiple applications

When combined with ultrahigh resolution fMRI, the BOLD signals elicited by neuronal activity at stimulated sites and connected sites can be detected and mapped. Compared to optogenetic stimulation, the advantage of INS-fMRI is its immediacy, permitting focal stimulation at any site in the brain in vivo without prior viral preparation. Although not cell type specific *per se*, the long range (e.g. cortico-cortical and cortico-subcortical) connections revealed necessarily arise from projecting pyramidal neurons (with exception of some gabaergic brainstem projections e.g. from amygdala also detected by INS-fMRI as evidenced by inverse intensity dependence). This method has broad applicability. It can be used at any brain site (e.g. cortical: V1^26^, V2^55^, SI^56^ or subcortical: amygdala^57-59^, pulvinar^60-61^), on different animal models (e.g. cat, squirrel monkey, macaque monkey), and on different MRI platforms (e.g. Siemens 7T, Varian 9.4T), and is compatible with or without contrast agents (MION)^26^. Possible applications include, for example, mapping high resolution connectomes^31^ (studying topography of connections^32,33^); identifying connected sites for targeting electrodes, anatomical tracers, or optogenetic viruses; studying feedforward vs feedback relationships between different nodes in the brain^26^ or studying development of columnar and laminar circuits within single individuals^62^. We have also developed methods (optical fiber bundles) for delivering sequences of focal INS stimuli to studying dynamics of circuit response as well as for brain-machine interface directions^15,63-64^. INS is also effective in evoking phosphones to which monkeys saccade^63^ and systematically modulating contrast discrimination behavior^65^, suggesting promise for simultaneous neuromodulation of circuits and behavior in ultrahigh field MRI.

### Need for broad dissemination

No other method offers this complement of capabilities *in vivo* and at brainwide scale. In addition, the fiber optic targeting and advancing method is accurate, inexpensive, and rapid. A published protocol would make this technique available so that it can be applied to different research questions. In particular, for the nonhuman primate research community (e.g. the Global Primate Data Exchange consortium, Prime-De^34^), this would provide a valuable additional tool for assessing neural circuits. Here we describe procedures for an application to generate largescale connectome data.

### Experimental design

The goal of the following procedures is to map the connections from focal stimulated sites in the brain using the INS-fMRI method. We provide an outline of the general strategy and specific implementation in Macaque monkey. These procedures can be adapted for other species, for different experimental aims, and can be implemented on different scanners. Thus far, these procedures have been used for both large (macaque) and small (squirrel monkeys) monkeys and cats; scanners have included Siemens 7T Magnetom and Varian 9.4T^26,32^. Below, we provide detailed description of the major steps in this protocol.

### Flowchart

INS-fMRI requires an MRI scanner (preferably ultrahigh field) and an 1870 – 1875nm laser. As shown in Figure 1, the primary steps are planning stimulation sites and, laser/fiber preparation, grid-chamber implantation, fiber insertion & targeting, anatomical and functional scanning during INS stimulation, and data analysis.

**Fig 1.**
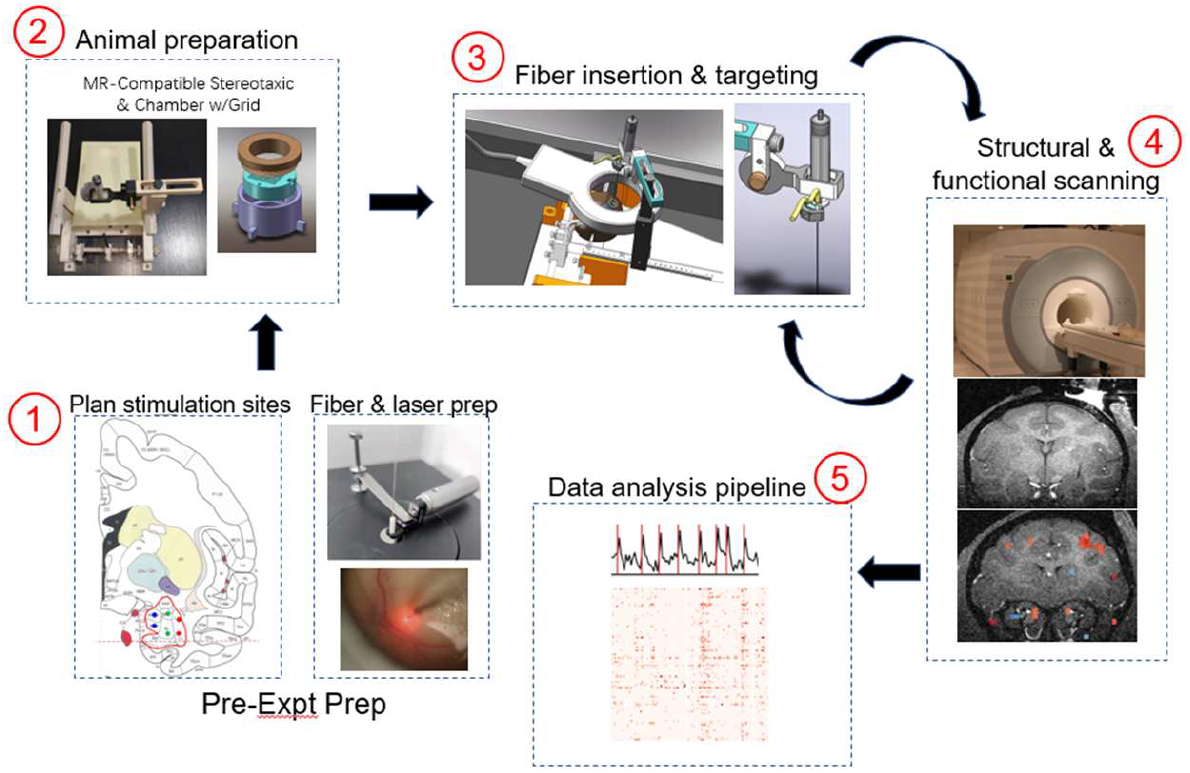
Flowchart of basic procedures for INS-fMRI. (1) Planning stimulation sites and preparation of optical fibers and laser. (2) Drilling burr holes in skull guided by grid. (3) Fiber insertion & accurate targeting. (4) Data collection: Structural & functional MRI scans. (5) Data analysis pipeline.

### ➀ Preparation of optical fiber probes

Optical fiber probes can be purchased cleaved to the desired length appropriate for the depth of the target site in the brain, e.g. for amygdala we chose 6cm long fibers attached to a 2.5mm diameter ferrule (ceramic to assure MRI compatibility). We use 200 µm diameter low-OH silica core fibers (0.22 NA) with 10 µm thick cladding of fluorine-doped silica and a 50 µm thick acrylic polymer coating, resulting in a total diameter (O.D.) of 320 µm. Prior to the experiment, the optical fibers need to be inspected, cleaned, and sterilized. Using a simple inverted microscope and appropriate optical mounting hardware, both the ferrule end and the free end of the fiber probes (the latter directly contacts the brain tissue) is inspected. If there are deep scratches or chips in the end of the fiber, discard the fiber. To remove any microscopic polymer coating chips from the core and cladding at the free end, the fiber may be drawn multiple times through a folded lens tissue wetted with isopropyl alcohol. Both ends must be free of contaminants such as dust or oil to achieve optimum transmission of the laser light. The fiber tip is cleaned by drawing the fiber tip across a folded sheet of lens paper wet with a drop of isopropyl alcohol from wet to dry area in one motion. Repeat this 2-3 times and reinspect to make sure the ends are clean.

### ➀ Calibration of laser power

The same laser current may produce different output intensities when optical fibers and connections are changed. Therefore, the laser output intensity at the fiber tip should be calibrated at the beginning of every experiment; this ensures that the cortex receives the expected stimulus intensity. To convert the target radiant exposure *RE* (0.1-1.0 J/cm^2^ per pulse) into laser intensity *LI*, use the following equation:

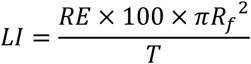

where the radius of the fiber core *Rf* is 100 µm (assuming that the illuminated area of the contacted brain tissue is equal to the cross-sectional area of the optical fiber) and the duration of a train of 100 pulses *T* is 0.5 sec. So, if the desired target radiation exposure is 0.1 J/cm^2^, the power detected is expected to be 6.3 mW. The power at the fiber tip can be measured using a power meter (Thorlabs, PM 100D, USA, Detector: Thorlabs S302C, USA), with the tip centered on the detector. The laser diode current is adjusted to achieve the target power output at the cannula tip. It is helpful to calibrate the laser current setting with the output power measured at the cannula tip and generate a lookup table that may be referred to during the experiment.

### ➀ Planning of stimulation sites and grid implantation

1. *Structural and vasculature brain scans (Figure 2A & 2B)*. Acquisition of high-quality structural scans (T1 image, voxels size 0.3 mm isotropic) is important for delineation of brain areas, alignment of the brain with standard maps (e.g. using AFNI, ITK-Snap, 3D slicer), and targeting desired sites. In addition, to minimize possibility of hitting major vessels, it is best to obtain a vascular scan.

*Preliminary plan of stimulation sites (Figure 2C, upper)*. Perhaps the most critical part of this experiment is planning the stimulation sites. It is helpful to know the data collection time per stimulation condition and the number of trials needed for getting significant signal (SNR) under anesthesia. If many points in a single region is desired, a grid can help to systematically, rapidly, and accurately acquire multiple points. The grid can be implanted in advance or on the experiment day.

### ➁ Chamber implantation with grid

1. *Plan of grid chamber implantation*. A grid, implanted at an appropriate location with a precise angle, offers ease and precision of access. To avoid vessels, we manually check each slice of structural images with planned penetration tracks as image overlays.
2. *Grid implantation surgery*. All procedures were carried out in accordance with NIH standards and with the approval of Zhejiang University Institutional Animal Care Committee. Following induction of anesthesia (for details^32^), the animal’s head is centered in the stereotaxic in the horizontal plane using a laser level projected on earbar to eyebar tip. The skull is exposed and fiduciary markers (two small drill holes or two crosses of tube filled with contrast agent lidocaine gel, Fig 3A1) are made on the skull. The animal is then moved into scanner to obtain an MPRAGE T1 scan. The animal is moved back to surgery room. The position and angle of the chamber implantation is determined (see Fig 3B) and the chamber fixed on the skull using dental cement. It is useful to hold the chamber with a micromanipulator at the desired angle during cementing. 2-3 ceramic screws are added to secure the chamber.
3. *Finalizing penetration tracks*. After chamber implantation, we plan the penetration tracks based on grid x, y and depth coordinates of target sites (Fig 3B). We make a MR mask file for these sites and highlight them on the acquired T1 and vessel maps. We then align our monkey brain with public atlas^35^ T1 images and parcellation maps by rigid body transformation (AFNI align_epi_anat.py) and elastic deformation (AFNI 3dQwarp) ^34^ of T1 images. From this procedure, a table of coordinates of the planned stimulation sites is generated.
4. *Burr hole drilling and fiber insertion*. The desired holes are marked on the skull by a fine pen tip inserted through each grid hole. The grid is removed and the skull holes are drilled (0.5mm hole size) at marked positions (Fig 5D).

**Figure 2.**
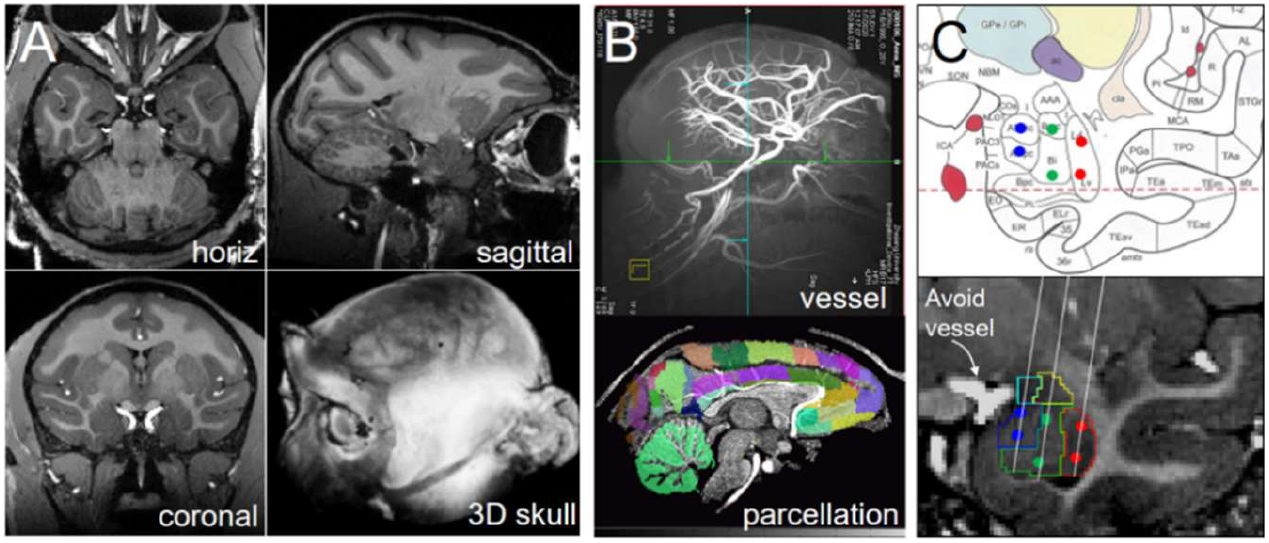
(A) Structural MPRAGE scans. (B) Vascular scan (top) and brain parcellation (bottom). (C) Stimulation sites (colored dots) on atlas of amygdala (top) and on structural MPRAGE scan. White lines: planned penetrations angled to avoid hitting large vessel (arrow)

**Figure 3.**
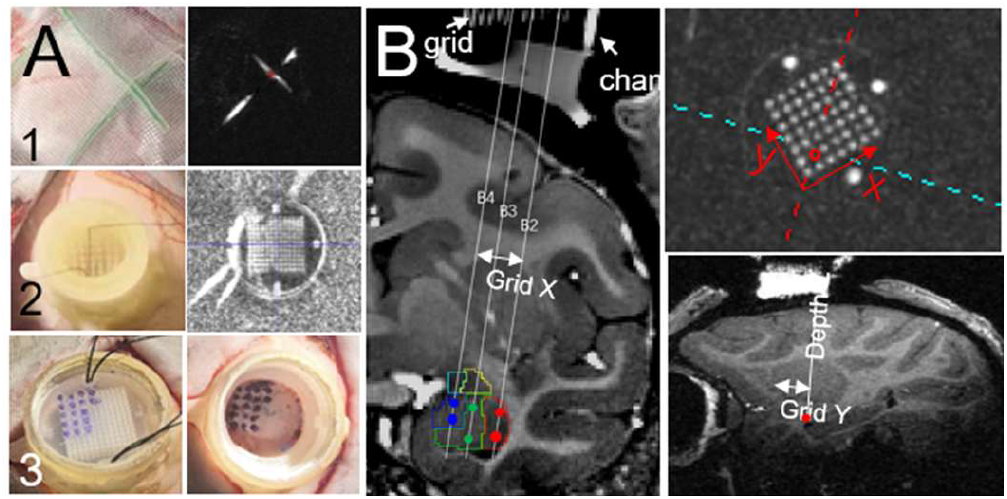
Grid implantation. (A) 1. Crossing of contrast agent tubes provide Fiduciary markers on skull (left) which appears in MPRAGE scan (right). 2. Insertion of grid in chamber (left); when filled indicator the grid holes appear white in MPRAGE (right). 3. Marking grid holes on skull (left). Thread used to remove grid. Drilling marked locations (right). (B) Structural images confirm chamber is in correct position for targeting desires sites (Grid X, Grid Y, depth Z).

### ➂ Moving animal between scanner and surgery room

Once the surgical procedures are completed, the animal, secured in the stereotaxic, is moved to the scanner. The head position must remain fixed in position in the stereotaxic. All support equipment in scanner must be ready to receive the animal (ventilator turned on, isoflurane turned on to 1%, and volume/rate set, ventilator tubes placed on the animal platform, SPO2/EKG mobile monitor ready, temperature pad pre-warmed). The animal’s endotracheal tube is then gently disconnected from the ventilator tubing with care not to dislodge endotracheal tube. Two persons lift the animal in the stereotaxic and quickly move it to the scanner bed. To prevent shifting of the brain in the skull, maintain the animal in horizontal position during the move. The stereotaxic is placed in a predetermined centered position on the scanner bed (head should be in center of the bore), the air tube is reconnected, monitor sensor attached to the animal, and vital signs (heartrate, SPO2, CO2, temperature) checked and adjustments made. The duration of air tube disconnection should not exceed 1 minute. IV lines are connected to an implanted catheter and saline infused for hydration (6-12 cc/kg/hr). All lines and tubes are taped securely to the bed to prevent pulling during movement of the bed into scanner. After completion of the experiment, these steps are reversed and the animal is returned to the surgery room for recovery.

### ➂ Fiber insertion and targeting

Before inserting the fiber, an ink mark is made on fiber probe 3∼4mm from the tip to indicate the approximate distance from the skull to the brain. A 26 gauge needle is similarly marked and manually inserted through the burr hole and the dura punctured. Some small damage of the underlying superficial cortical layers may occur. Then the fiber optic probe is carefully inserted by hand through the burr hole and held in position while a second person attaches the ferrule end of the probe to an MRI-compatible 3D-printed holder (see Equipment section) (Figure A, C). Center the fiber tip in the burr hole so that the fiber does not rub against the wall of the burr hole, which may lead to a curved trajectory. Once the tip is in the brain, the fiber optic probe is then inserted to a predetermined depth using an MR-compatible hydraulic fiber pusher (Figure 4B).

**Figure 4.**
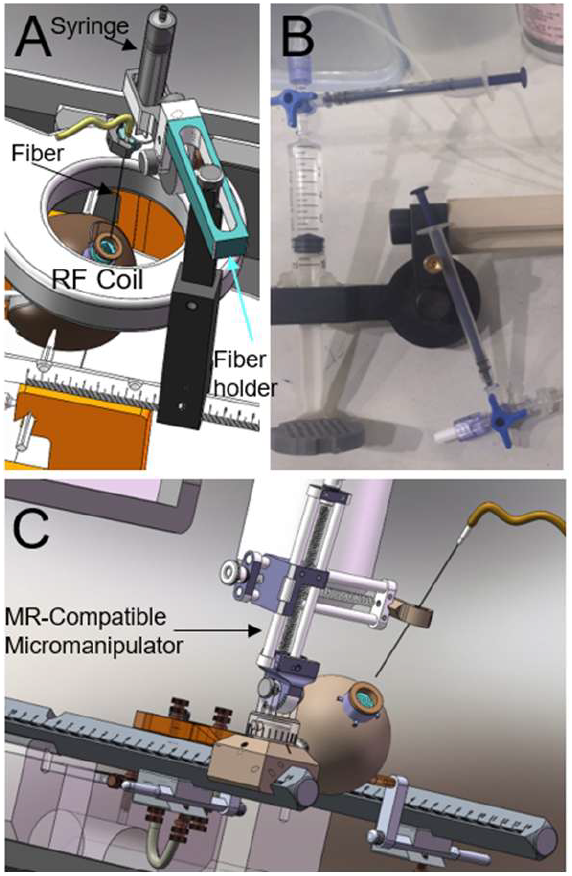
Fiber positioning and pushing. (A) Rendition of setup. (B) Fiber pushing system composed of large and small syringes connected by a T-valve, and tubing. Pushing of fiber into brain is done from control room. (C) MR-compatible micromanipulator for flexible positioning of fiber into burr hole.

**Figure 5.**
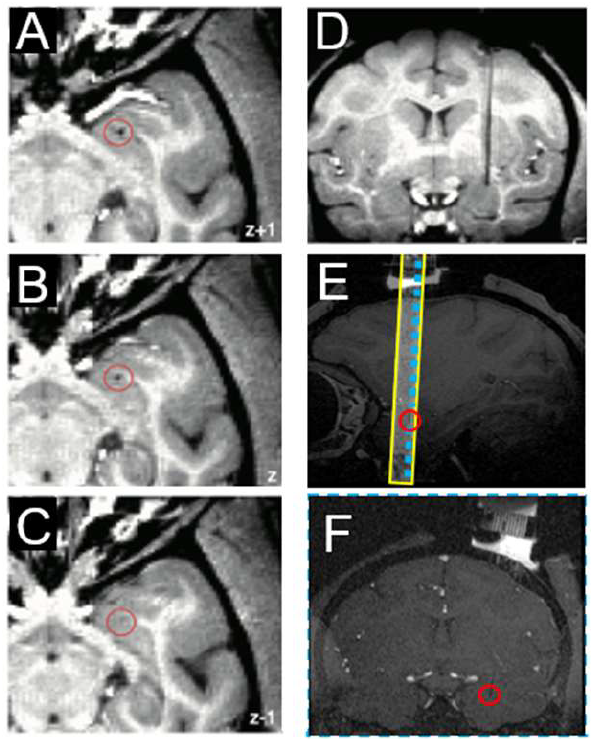
Locating fiber tip. (A-C) Three sequential horizontal sections from dorsal to ventral. Fiber tip (dark dot in red circle) is visible in first two sections, but not in third. Depth of tip is the most ventral horizontal section where dark dot is still visible. (D) Coronal section reveals AP and ML coordinates of fiber tip. (E) Fast structural scan acquires only a few slices (yellow box on saggital section). (F) Blue dashed line: slice

### ➃ Acquisition of MR images

MR images were acquired on a 7-T research scanner (Siemens Health-care, Erlangen, Germany) equipped with a whole-body gradient set (70 mT/m and 200 T/m per second). The specific location of the fiber tip is then confirmed either via an MPRAGE scan (Figure 5A-D) or a fast structural scan (FLASH sequence^36^, T1 image, voxel size 0.3 mm, 3-4 slices near the fiber tip) (Fig54E-F). The partial brain FLASH has low gray/white matter contrast, but it saves valuable experimental time. Our strategy for aligning the FLASH image to reference anatomical image is to first use the MPRAGE image for alignment and then apply the alignment transformation to the FLASH image to confirm tip location. Note that there are times when the pusher may not advance the desired amount. Hence, documentation of precise tip location is critical for interpretation of functional stimulation data. To further increase SNR, a single loop RF coil (RAPID MR International, Columbus, OH, USA) for was securely positioned above the animal’s head (Figure 4A). Any motion or vibration of the coil will introduce noise and degrade image quality. This coil provides improved homogeneity of temporal signal-to-noise ratio (tSNR) over regular surface coils, resulting in images with similar tSNR values (mean tSNR of gray matter ∼75). Functional images from opposite phase-encoding direction are also acquired for correction of image distortion^37^.

INS stimulation trials are then conducted at various depths along a single penetration pathway. In each trial, functional images of BOLD signals were obtained with a single-shot EPI sequence at a voxel size of 1.0-mm isotropic (50 slices) or 1.5-mm isotropic (35 slices) (TE, 22 ms; TR, 2000 ms; matrix size, 64 × 64; flip angle, 90°). MR acquisition is synchronized with laser stimulation onset via a pulse generator (AMPI Master 9, Israel) controlled using MATLAB. Upon completion of stimulations, the fiber is slowly withdrawn using the fiber pusher. This procedure is then repeated in another burr hole.

### ➃ Stimulation paradigm

INS (infrared neural stimulation) was carried out using a near-infrared diode laser (Changchun New Industries, FC-W-1870) at 1870±20nm wavelength guided through a 200-μm diameter fiber (0.22 numerical aperture). The basic INS train is 0.5 s in duration consisting of 250 μs wide pulses delivered at a repetition frequency of 200 Hz (i.e., a train of 100 pulses) (Fig 6A). Radiant exposures ranged from 0.1–1.0 (J/cm^2^)^14,13,38^.

**Figure 6.**
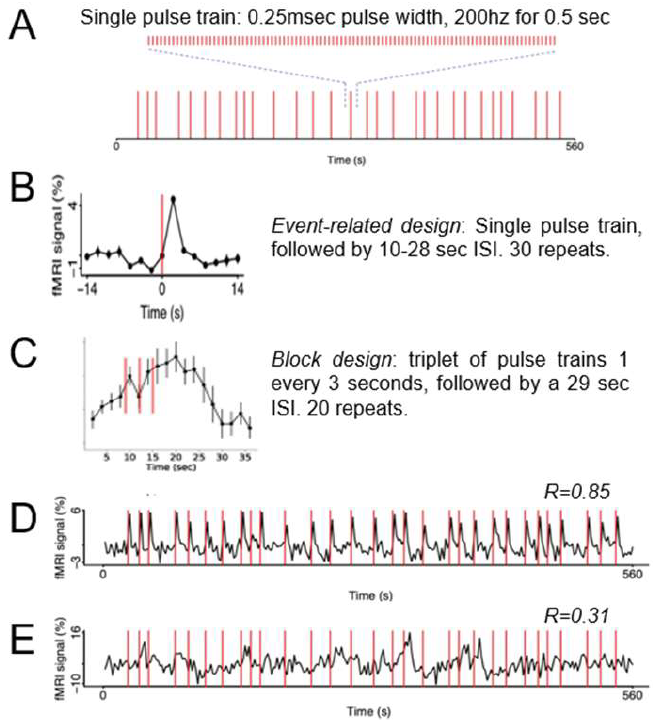
INS stimulation. Single pulse train (A) delivered either in event-related (B) or block design (C). Stimulation can result in high (D) or low (E) correlation.

**Fig. 7.**
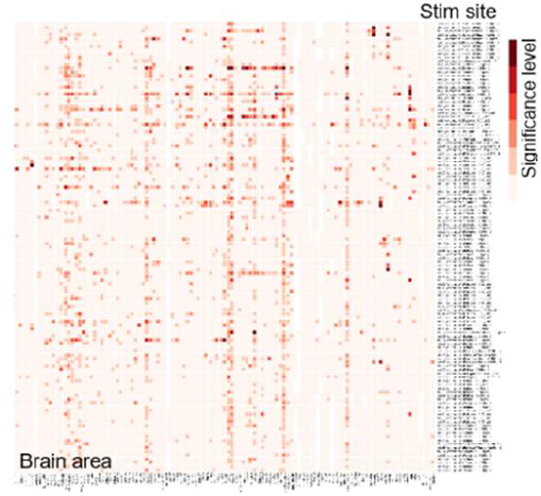
Connectome of all stimulated sites in a single brain area with all other areas of the brain. (Roe et al, Organization for Human Brain Mapping Abstract 2021)

Either event-related (Fig 6B) or block design (Fig 6C) paradigm may be used^32,26^. Event-related design provides better understanding of the BOLD hemodynamic response function induced by INS; block design is more robust and offers better SNR. We conduct at least two runs for each stimulation site at each INS intensity to improve SNR and to test reproducibility of results. Typically, data from 2-4 intensities (i.e., 0.1, 0.2, 0.3J/cm^2^ per pulse) are acquired at each site.

Effective intensities used for cortical surface sites with and without artificial dura, and for subcortical sites may differ.

### ➃ Animal procedures & anesthesia

A good anesthetic plane is important for establishing a stable baseline for acquiring functional images. In our preparation, this was achieved using sufentanil (IV infusion, induction: 0.5μg/kg, maintenance: 1μg/(kg*h)), supplemented with up to 0.3% isoflurane. Vital signs were monitored including heart rate, CO2, and temperature. Drugs were delivered in saline or 5% dextrose continuous IV infusion. Each session lasts no more than 24 hours. The animal is recovered and typically resumes normal activity and appetite the next day. and sessions are spaced 1-2 weeks apart to allow the animal sufficient recovery time. The optical fibers typically do not leave lasting evidence of tissue damage: by the next session, no tracks are seen in the online brain scans. Careful planning of the fiber trajectory can minimize the chance of puncturing large blood vessels. However, in the case that a vessel is damaged, bleeding may cause brain swelling. Should an animal exhibit discomfort, analgesics are provided for pain and dexamethasone or a diuretic provided to reduce edema. Care and treatment is supervised in consultation with a nonhuman primate veterinarian. Although rare, in cases where the animal’s health is compromised, the animal may be removed from study or euthanized.

### ➃ Documentation

Systematic documentation is important. Experimental notes are saved as a digital document including both text and pictures. It is organized as lines of events with time stamps. The events include the surgery, fiber location, vital sign monitoring, laser stimulation parameters, and MR scans.

### ➄ Data analysis: Characterization of stimulation sites

To characterize the sites that we actually stimulated, we align all the structural images of fiber tips together in the same brain space. We then manually determine the fiber tip locations and write down their coordinates. The alignment is done by AFNI program and rigid body transformation. The fiber tip locations are determined by checking each horizontal slice and finding the slice in which the optical fiber artifact disappears (Fig 5). The aligned structural images are averaged together to create a high-contrast and low-noise structural image. Coordinates of fiber tip locations (stimulation sites) are then marked on the average structural image with in-house programs. Stimulation sites are then assigned to different brain areas using atlas parcellation. A summary table of stimulation sites and connected sites is produced.

### ➄ Data analysis: Activation maps and connection detection

Using standard fMRI analysis, connected sites are determined by identifying significant correlation between the simulated site and each voxel in the brain (Fig 6D-E). BOLD functional images are converted to NIfTI and AFNI format, preprocessed with correction for slice timing, motion, image distortion and baseline shift. Significant responses are identified in a commonly used generalized linear model (GLM) approach, in which the time-course of each voxel is regressed on the stimulus predictor. Regression coefficients are subjected to T-tests and multiple testing correction with Benjamini-Hochberg method. Voxels with significant T-test values are overlain on structural images. Individual voxel time-courses are extracted from EPI data, then averaged across trials, and plotted. Each baseline is estimated with the mean MR signal over the full time-course. The analyses are done with software AFNI, Nipype and in-house programs written in Bash, Python, and R.

### ➄ Data analysis: Robustness tests on detected connections

Both permutation test and half-data analysis are carried out to examine the robustness of identified connections^32^. The permutation test is complementary to the multiple testing correction done using the Benjamini-Hochberg method. In the permutation test, we randomly shuffle the functional time-courses of voxels to create a large number (usually 1000) of randomized data sets. We then apply the same fMRI analysis on these randomized data sets and compare the result to that from the real data. For each voxel, the permutation p-value is defined as the proportion of randomized time-courses that are more or equally significant than the real time-course. This permutation test p-value represents the significance of the detected response.

The half-data analysis tests for reproducibility. In the half-data analysis, the full time-course of each voxel is divided into odd and even trials. This results in two halves of the data subjected separately to GLM tests. This also results in two sets of activation maps compared side by side. The significant voxels common to the two activation maps are considered as reproducible.

## MATERIALS

### Facility

MRI system (Siemens, Erlangen, Germany) Small surgical procedure room near MRI

### Equipment

MR-compatible stereotaxic compatible with vision (custom-made, PEEK) MR compatible ear bars & eye bars (custom-made, PEEK and copper) MR-compatible micromanipulator (custom-made, PEEK) Leveling Laser on tripod (German, BOSCH, GLL5-50X) Vital signs monitor (China, Mindray, IPM12 Vet) MR compatible pulse oximetry & EKG sensor (USA, SA Instruments, Model 1030) Micro syringe pumps (WZ-50C6) Ventilator (Italia, UGO 6125-100) Anesthesia apparatus (China, RWD, R580) Water thermal system (PolyScience, PD28R-30-A12E) RF Coil (RAPID MR International, Columbus, OH, USA) Infrared Diode Laser 1870±20nm (China, Changchun New Industries, FC-W-1870) Pulse Generator (Israel, AMPI, Master 9) Fiber Patch Cables: 10 m long, 200 µm core diameter (0.22NA) with terminations appropriate for connection to laser and fiber probe.

Fiber Probe: 6.5cm long, 200 µm core diameter (0.22NA); outer diameter 320um, mounted in a 2.5mm ceramic ferrule.

Power meter and detector (USA, Thorlabs PM 100D with S302C detector)

### Drugs

sufentanil (china, renfuyaoye, SFDAH20054171)

Isoflurane, maintain for scan 0.2%, (china, RWD, R510-22)

Atropine intramuscular, (Mkdoctor, syz070011510)

Sterilized saline, (china, guangxiyuyuan, SFDAH45020976)

Sterilized 5% dextrose, (china, guangxiyuyuan, SFDAH45020562)

Zotetil50 (virbac china, wsyz43)

Pentobarbital (Sigma-Aldrich, 57-33-0)

Paraformaldehyde,PFA (Sigma-Aldrich, P6148-1KG)

### Surgery tools

Scalpel (China, shanghaijinzhong)

surgical scissors (China, shanghaijinzhong)

forceps (China, shanghaijinzhong)

hemostatic forceps (China, shanghaijinzhong)

osteotome (China, shanghaijinzhong)

dental engine (China, Strong)

dental cement filler (China, shanghaijinzhong)

ceramic bone screw (USA, Thomas recording, AN000055)

ceramic bone screw driver (USA, Thomas recording)

Surgical Suture (China, shanghaijinhuan)

## EQUIPMENT SETUP

### Synchronization between MR scanner and laser

The MRI scanner console and the pulse generator are connected to the stimulus control computer (SC-PC) via USB port. Once the MR scanner initiates image acquisition, a trigger is sent to SC-PC, which starts generating trigger signals for pulse generator according to customized paradigm program. Pulse generator enable each trigger signal to start a pulse train for laser. Pulse number, width and duty cycle of the train are preset in pulse generator. A long fiber optic runs from the laser to the animal in the scanner. This permits the synchronization between laser stimulation and MR data acquisition.

### MR-compatible micromanipulator and fiber pushing system

Our MR-compatible micromanipulator used for positioning the fiber is modelled on a Kopf micromanipulator and is made from peek and the turning screw from bronze (see Fig 4C). To push the fiber probe into the brain remotely, we use a water-based hydraulic system comprising an injection syringe with extension tube and T-junction (Fig 4B). A 10 ml syringe is connected to the ferrule on the optic fiber probe using a 3D-printed holder. Another 1ml syringe connects to the 10ml syringe by an extension tube. This is then calibrated to translate milliliters of fluid into millimeters of fiber tip movement in the brain. To achieve accurate movement of fiber tip, it is important to remove all air from this system and for connections to be tight.

## PROCEDURE

### Planning stimulation sites and penetrations

1 Obtain high-resolution structural image of the whole monkey brain (T1 image, voxel size 0.3 mm isotropic) with head secured in MR-compatible stereotaxic in the eye bar/ear bar defined horizontal plane.
2 Align digital monkey atlas (D99)^35^ to our monkey brain with AFNI programs. Identify target brain areas with the aligned atlas parcellations.
3 Plan penetration tracks (number of stimulation sites, locations, avoid vessels). If needed, plan chamber implantation (by determining stereotaxic coordinates and angle of the grid in chamber).
4 Make a preliminary plan of stimulation sites.

### Optional: Installation of grid chamber (Timing 3-4 h)

5 Remove food from the monkey at least 12 hours before experiment.
6 Inject atropine (0.03mg/kg intramuscular, IM) to reduce secretions. Inject Zoletil50 (2.5mg/kg IM) for sedation. Wait about 5-15minutes for drugs to take effect.
7 Intubate animal with appropriate endotracheal tube. Implant intravenous (IV) catheters: place 2 IV catheters, one in each saphenous vein.
8 Place monkey head in the stereotaxic apparatus so that eye bars are in orbit of eye and ear bars are symmetric. Make sure the head is in the middle of the stereotaxis. Make sure the earhole and eye orbit at the same horizontal plane. CRITICAL STEP: use a horizontal laser level to make sure that the earhole and the lower eye socket are at the same level.
9 Remove the hair on head by depilatory cream, then sterilize skin by lidocaine and 75% alcohol each 3 times. Inject the lidocaine of the surgery area to reduce pain.
10 Open the skin and expose the skull, remove muscles if needed. Then clean the surface of the skull by hydrogen peroxide to move all tissues.
11 Drill 2 tiny holes on the skull as reference points (Better not to penetrate the skull). Inject lidocaine gel into the sockets.
12 Move the animal inside the MRI bore for a high-resolution T1 image (voxel size 0.3 mm isotropic).
13 Identify targeted brain area. Make a final plan of grid implantation. Record the stereotaxic coordinates and angle of the grid center. Also calculate the distances between the reference points and the grid center.
14 Move the animal to the surgery room. Use a micromanipulator to locate the planned grid centers by using both the stereotaxic coordinates and the reference points. Check
15 if these two ways of localization agree with each other.
16 Install grid chamber on the skull at the planned location and angle. Fix it with dental cement and ceramic screws.
17 Insert the grid inside the grid chamber. Fill the grid and chamber with MR contrast agent gadolinium. Use a threaded ring to secure the grid inside the chamber.
18 CRITICAL STEP: To image clearly all the grid holes, inject enough gadolinium into all the holes in the grid and avoid air bubbles.
19 Put animal inside MRI bore. Get structural images with both the brain and the grid (T1 images, voxel size 0.3 mm isotropic).
20 Adjust the plan of stimulation sites and penetration tracks. Determine the information for each site (site number, brain area, name of the grid hole, depth).
21 Move the animal to surgery room for burr hole drilling. If stimulation and imaging will be done on another day, recover the animal.

### Stimulation and imaging (can be repeated multiple times in a single animal)

22 Mark the chosen holes on the skull with a fine surgery pen through the grid holes. Use a fine drill bit (diameter 0.4 mm) to drill the holes in the skull. Clean the skull surface after drilling.
23 Move the animal to scanner bed. Make sure the animal is in a stable anesthetized state.
24 Confirm that the fiber cable is correctly connected to the laser. Test the visible guide laser output (usually red light).
25 Use a fine stainless steel syringe needle (diameter 0.3-0.4 mm) to poke the dura (it is ok to bring this needle into scanner room).
26 Mark the required depth for first insertion on fiber optic probe.. Place fiber probe in the MR-compatible fiber holder. Move the fiber tip into the chosen grid hole and corresponding skull hole while keeping it straight. Use the syringe pusher to push the fiber inside the brain until the marked line on the fiber reaches the grid surface. This ensure the fiber tip into brain tissue and thus visible in imaging.
27 Obtain a whole brain structural image (T1 image with MPRAGE sequence, voxel size 0.3 mm isotropic). Confirm the depth of the fiber tip in the structural images and record.
28 If the fiber tip has not reached expected depth, use the fiber pusher to move the fiber again. Avoid puling the fiber backward,, as it may not return reliably to the expected position. Find the MR coordinates of the fiber tip and the angle of the coronal slice containing the whole fiber inside the brain. Use these coordinate and angle information to get a structural image of the few coronal slices surrounding the fiber with the FLASH sequence (T1 image, voxel size 0.3 mm).
29 Once fiber tip reaches the target site, begin laser stimulation trials and acquisition of fMRI images.
30 Repeat steps 26 and 27 for desired sites along this penetration trajectory. Repeat steps 24-27 for sites along another penetration trajectory via another grid hole.

### Animal recovery

31 The animal is recovered after data collection. Move animal to surgery room, with saline and glucose maintain as 5ml/h and also isoflurane to 1.5∼2%. Clean the skull and skin using hydrogen peroxide 3 times. Remove grid and rinse inside of chamber with saline with 5% ceftriaxone sodium until clean. Place a small piece of gauze with drops of amikacin, close the chamber with a cap, and use bone wax to seal the gap between cap and chamber. Give atropine (i.m. 0.06mgy/kg, to reduce mucous secretions), narcon (0.06mg/kg, sufentanil reversal agent), ceftriaxone sodium 25mg/kg (i.m. antibiotic). Extubate when animal can breathe stably and maintain appropriate CO2 levels. Return to cage and monitor closely until fully alert. **Timing: 1h**.

### Data processing

32 Characterize the stimulation sites with structural images containing the artifacts showing the fiber tips. Select the sites associated with brain areas of interest.
33 Obtain activation maps of all functional scans. Identify connections while controlling the rate of false positives at or below 5%.
34 Construct connectivity matrix based on the best voxel response in each brain area.
35 Perform additional robustness tests on the detected connections.

### Timing

Planning: hours, days

Chamber Implantation procedure: 3-4 hours

Data acquisition: 30-60 minutes per site, up to 18 hours for recovery session, 1-4 days for terminal experiment.

Depending on the length of sessions, sessions can be repeated once every 1-2 weeks.

**Table 1:**
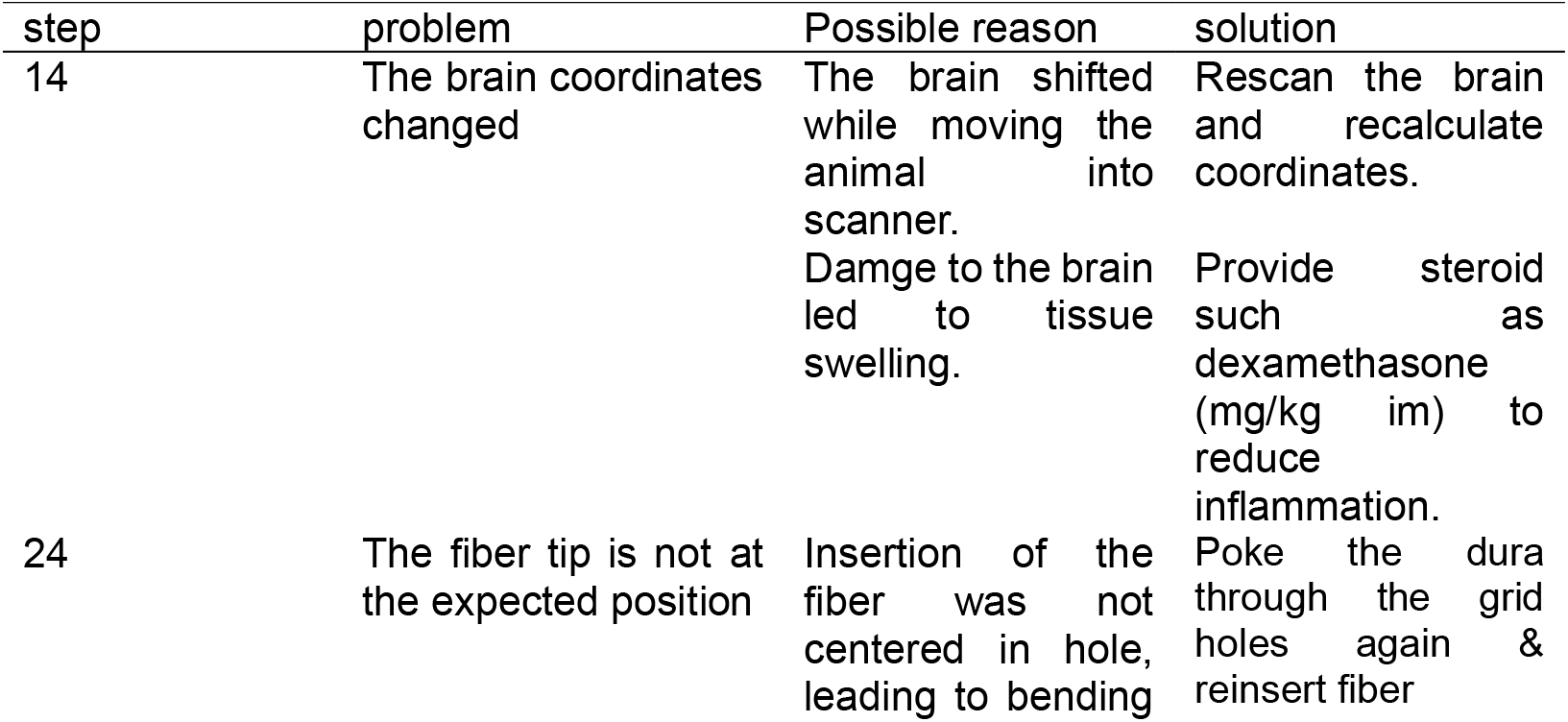
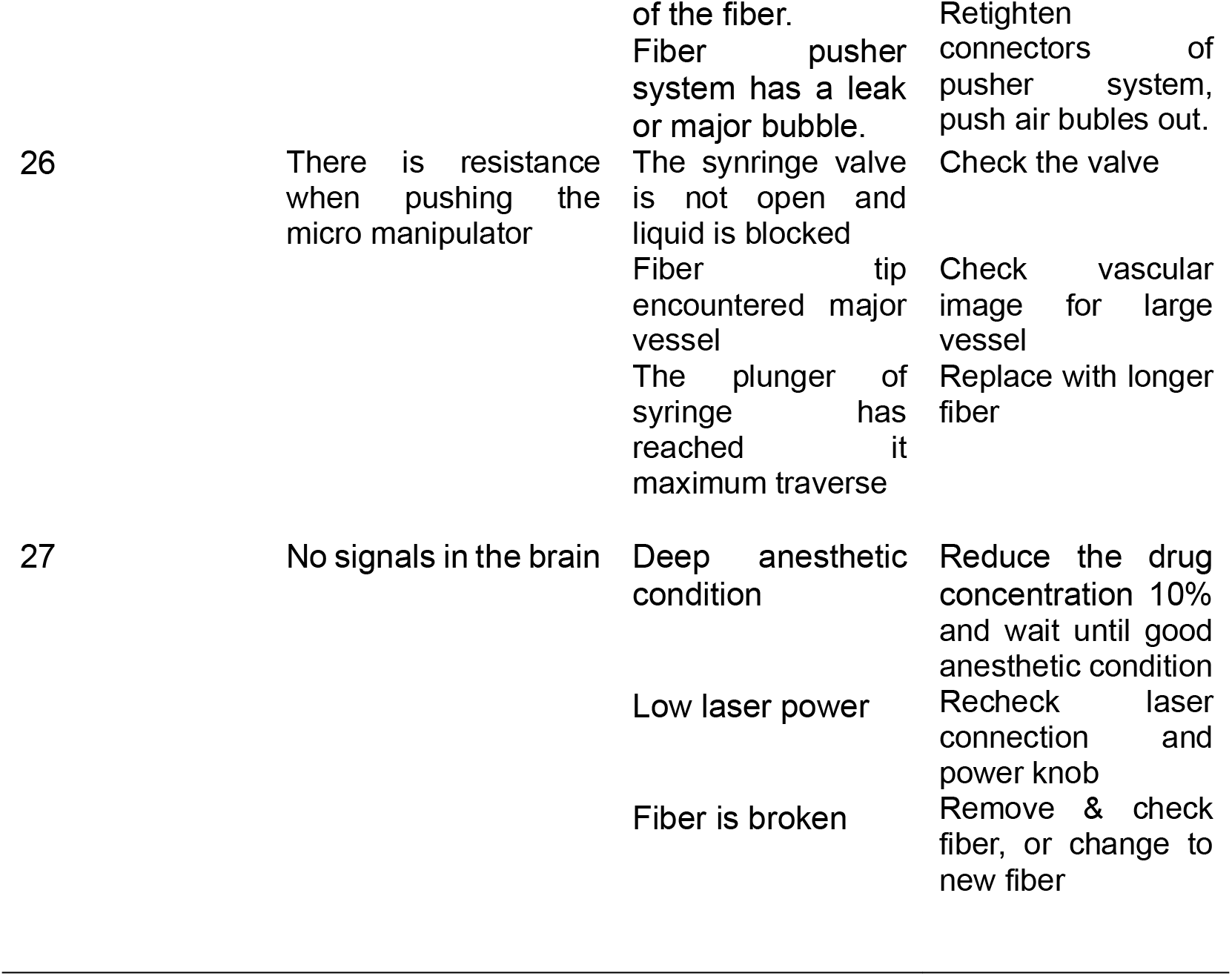
Troubleshooting.

### Troubleshooting

*Troubleshooting 27: Online assessment during experiment:* In order to assess whether the INS is effectively inducing brain activation during the experiment, we conduct online analysis to check the GLM statistical results after each run. The functional activation map and time courses are quickly assessed to visualize statistically significant voxels (examples shown in Fig D & E). Roughly, if the response onset time courses correlate well with the stimulation onset times and signal change magnitude is reasonable (e.g. > 2%), the activation is deemed acceptable. This provides some feedback regarding whether the animal is at a good level of anesthesia (too deep: no neural signal, too light: increased baseline variability) and whether the laser tip is delivering stimulation. If significant signal is not obtained, troubleshooting should include examination of anesthesia dosage (recommend adjust by 10%), checking the body temperature (low temperature can drastically reduce signal), and confirmation that the laser is functioning properly.

*Anesthesia:* For any experiment in anesthetized animals, it is important to maintain a stable anesthetic plane. Overly deep plane will lead to loss of neuronal response; a plane that is too light results in fluctuating baseline and results in poor signal-to-noise. We have accumulated experience with different anesthetics in monkeys and find that sufentanil with the described induction and maintenance doses provides a reasonably stable baseline. Isoflurane is used as a secondary ‘tweaker’ as it its effects are rapid and reversible; isoflurane is known to suppress neural signals, so we typically do not use more than 0.4% in conjunction with sufentanil.

*Animal Monitoring:* As our respirator is outside the scanner room, the volume within the ventilator tubes must be taken into account. Testing respiration system with a balloon of known volume is recommended. Similarly, for temperature maintenance, a large reservoir water bath is needed and tubes need to be insulated to minimize heat loss. Monitors must be tested rigorously to ensure that heartrate, C02 sensor, SP02, and temperature readings are reliable. I.V. lines are long and flow must be maintained to prevent clotting; lines must be checked for air bubbles and leaks at connectors.

## Anticipated Results

### ➄ Visualization of data

*Brain activation maps*. For each run, the significant voxels are rendered on a series of brain slices (e.g. see^32^ Fig 4). Multiple significance levels may be visually inspected (e.g. p<0.001, p<0.0001) to guide further analysis strategies.

*Connectivity matrix*. After stimulation and detection of connections, we construct a connectivity matrix between stimulation sites and all known areas in a monkey brain. This needs to be done in several steps. Significant voxels are labeled with brain area names. Then each known brain area is represented by the most significant p-value of all voxels in that area. Lastly, minus log10 p-values are calculated and plotted in a heatmap for visualization. This whole procedure is done with AFNI programs and in-house written programs.

### Evaluation of the Method

*Mechanism underlying INS*. Unlike optogenetics, INS is not specific to cell type. It is likely to affect multiple components in the parenchyma, including neurons, glia, and vasculature, and may therefore have other effects beyond neural activation. The mechanism underlying INS is still unclear. Evidence suggests that heat transients induced by the brief INS pulses leads to changes in membrane capacitance and concomitant Na and K flux, which in turn produce neural activation^28-30^. Other possible mechanisms include activation of heat-sensitive TRP receptors^39,40^ and potential mechanical effects ^41,42^ of optical stimulation; however, these possibilities are considered less likely in the primate brain. Despite this, there have been several exciting applications of INS, including cardiac peacemaking^43^, cochlear stimulation^44^, sciatic nerve stimulation^42^, inhibition of pain perception^45^, and suppression of action potential generation at neuromuscular junctions^30^.

*Assessing the data:* (1) *First & second synapse*. In addition to assessing statistical significance and reliability of the data, another important assessment is whether significant voxels are due to direct or indirect connections. Development of methodology to distinguish direct vs indirect connections is still in progress. In our experience, the focal and brief nature of the stimulation leads to robust activation of the first synapse and a smaller activation (e.g. by 10-50% of first synapse amplitude) and greater variability at the second synapse (presumed to be non-direct from known anatomical studies^26^). This predicts that third synapse activations are unlikely to reach significance. Our comparisons of activation patterns to anatomy are also consistent with first and second synapse activation^26,32^. (2) *Orthodromic vs antidromic*. Consistent with electrical stimulation studies^8^ (e.g. Messinger et al 2015 SFN abstract), focal INS stimulation leads to activation patterns more consistent with orthodromic activation^26^. This has been inferred from comparisons with non-reciprocal anatomical connections (e.g. area 2 projects in feedforward fashion to areas M1 and 3a and feedback fashion to area 3b and 1 ^26^; our amygdala stimulations also replicate Messinger et al 2015’s findings suggesting effects are orthodromic). (3) *Suppression*: We consistently observe that, while we do observe intensity dependence within a certain range, stimulation at high intensities leads to fewer activated voxels. This has also been observed with electrical stimulation^20,46^. Thus, at least at this stage, we tend to focus on the lower stimulation levels, which may have more limited contribution from complex polysynaptic effects. Note that it is important to remain below damage threshholds^47^.

*Future development*. INS is developing into a viable option for focal domain-specific brain stmulation. In combination with other methods such as fMRI, it provides additional options for studying brain circuits in vivo and at whole brain scale.

## Acknowledgements

This work was supported by: National Key R&D Program of China 2018YFA0701400 (to A.W.R.), Chinese NSF Instrumentation Grant No. 31627802 (A.W.R.), Key Research and Development Program of Zhejiang Province 2020C03004 (to A.W.R.), Fundamental Research Funds for the Central Universities 2019XZZX003-20 (to A.W.R.), NSFC 81961128029 (to A.W.R.)

